# TMEM55B regulates lysosomal acidification and ER–lysosome calcium signaling to promote lipolysis

**DOI:** 10.1101/2025.10.21.683636

**Authors:** Nagi Mukae, Honoka Maki, Michiko Shirane

## Abstract

The lipid transfer protein PDZD8 promotes cholesterol metabolism by facilitating endosomal maturation at membrane contact sites (MCSs) between the endoplasmic reticulum (ER) and endolysosomal compartments. Endosomal maturation is closely associated with lysosomal acidification and calcium signaling; however, the molecular mechanisms linking PDZD8 to these processes remain unclear. Here, we identified TMEM55B as a lysosomal PDZD8-associated protein and a candidate regulator of cholesterol metabolism. TMEM55B has been reported to interact with the v-ATPase complex, suggesting a role in lysosomal function. Suppression of TMEM55B expression reduced lysosomal acidification and impaired lipid droplet turnover. In addition, TMEM55B depletion attenuated lysosomal Ca^2+^ release and reuptake and diminished ATP-induced Ca^2+^ responses in the ER, consistent with impaired calcium-induced calcium release (CICR) at ER–lysosome MCSs. By contrast, mitochondrial Ca^2+^ dynamics were unaffected. These findings identify TMEM55B as a regulator of lysosomal acidification and calcium dynamics and suggest that the PDZD8–TMEM55B axis promotes lipolysis and cholesterol metabolism through coordinated regulation of lysosomal function at ER–lysosome membrane contact sites.

## Introduction

Membrane contact sites (MCSs) between the endoplasmic reticulum (ER) and late endosomes/lysosomes (LE/Ly) are established through specific tethering complexes rather than random membrane encounters. ORP1L and VAP were among the first proteins identified as ER–LE/Ly tethering factors, and subsequent studies have shown that numerous lipid transfer and trafficking proteins—including members of the ORP, STARD, and LAM families, as well as NPC1, Protrudin, and PDZD8—function at these sites^1-3^. ER–LE/Ly contacts increase during endosomal maturation, a process closely associated with both lysosomal acidification and Ca^2+^ signaling. Cholesterol and Ca^2+^ therefore serve as key mediators of communication between these organelles at ER– LE/Ly MCSs^4^.

Cells obtain cholesterol through two major pathways: de novo synthesis and endocytic uptake of lipoproteins (LPs) (LPs)^5-7^. In the brain, cholesterol is produced predominantly by glial cells and delivered to neurons via HDL-like lipoproteins. Following receptor-mediated endocytosis, cholesteryl esters (CEs) are hydrolyzed by lysosomal acid lipases to generate free cholesterol, which is subsequently distributed to the ER and other organelles for cellular functions^4,7-9^. In parallel, CEs stored in lipid droplets (LDs) can be degraded through lysosome-dependent lipolysis, allowing lysosomal lipases to mobilize stored lipids^10-13^. Efficient lysosomal maturation and acidification are therefore essential for cholesterol metabolism derived from both LPs and LDs.

PDZD8 is an ER membrane protein originally identified as a Protrudin-interacting protein and functions at ER–LE/Ly MCSs^14-20^. PDZD8 belongs to the TULIP superfamily and contains a synaptotagmin-like mitochondrial lipid-binding protein (SMP) domain that mediates lipid transfer^16,21,22^. Together with Protrudin and VAP proteins, PDZD8 contributes to cholesterol trafficking at ER–LE/Ly MCSs^3^. We previously demonstrated that PDZD8 deficiency impairs lipolysis and causes CE accumulation in the brain, indicating an important role for PDZD8 in cholesterol metabolism^23-25^. Consistent with these findings, abnormal CE accumulation has also been reported in Alzheimer’s disease (AD) patients and AD mouse models^26-28^. Moreover, neurons carrying the APOE4 allele, the strongest genetic risk factor for AD, exhibit lysosomal dysfunction accompanied by CE accumulation and lipofuscin formation^29,30^. Despite the established role of PDZD8 in cholesterol metabolism, the mechanisms by which it regulates lysosomal function remain poorly understood.

In addition to cholesterol metabolism, increasing attention has focused on lysosomal Ca^2+^ signaling. During endosomal maturation, luminal acidification is accompanied by progressive accumulation of Ca^2+^ within endolysosomal compartments, making lysosomes an important intracellular Ca^2+^ store^31-34^. Luminal Ca^2+^ promotes endosomal maturation through intraluminal vesicle fusion, whereas Ca^2+^ release from lysosomes contributes to diverse signaling pathways. In particular, calcium-induced calcium release (CICR), whereby lysosomal Ca^2+^ release triggers secondary Ca^2+^ release from ER-localized IP3 receptors and ryanodine receptors, has emerged as an important mechanism for intracellular Ca^2+^ signaling^35-39^. Endolysosomal acidification also regulates the activity of lysosomal Ca^2+^ channels such as two-pore channel 2 (TPC2) and TRPML1. Although Ca^2+^ signaling at ER–LE/Ly MCSs has been implicated in multiple cellular processes^34,40^, the molecular mechanisms coordinating lysosomal acidification, Ca^2+^ dynamics, and lipid metabolism remain incompletely understood.

TMEM55B is a lysosomal membrane protein with multiple reported functions. Originally identified as a phosphatidylinositol phosphatase that converts PtdIns(4,5)P_2_ to PtdIns(5)P^41,42^, TMEM55B was subsequently shown to associate with the v-ATPase and Ragulator complexes, thereby contributing to lysosomal signaling and mTORC1 activation^43^. Consistent with its importance, TMEM55B knockout mice are embryonic lethal^43^. Because v-ATPase is essential for endolysosomal acidification, disruption of this pathway results in lysosomal dysfunction and storage disorders^44-46^. TMEM55B has also been implicated in dynein-dependent retrograde lysosomal transport, oxidative stress responses, and regulation of LDL receptor expression^47-50^. However, whether TMEM55B contributes to lipid metabolism through regulation of lysosomal acidification and Ca^2+^ signaling remains unknown.

In the present study, we identify TMEM55B as a PDZD8-associated lysosomal protein and demonstrate that TMEM55B regulates lysosomal acidification, lysosomal and ER Ca^2+^ dynamics, and lipid droplet turnover. Our findings reveal a previously unrecognized PDZD8–TMEM55B pathway that links lysosomal function to lipolysis and cholesterol metabolism at ER–LE/Ly MCSs.

## Results

### PDZD8 interacts with the lysosomal protein TMEM55B

In neuronal cells, the lipid transfer protein PDZD8 promotes cholesterol metabolism via endosomal maturation; however, the mechanism by which this ER protein regulates lysosomal activity remains unclear. To identify lysosomal proteins interacting with PDZD8, we established a Neuro2A cell line stably expressing 3×FLAG-tagged PDZD8. The PDZD8 complex was isolated by anti-FLAG immunoprecipitation, separated by SDS-PAGE, digested with trypsin, and analyzed by LC-MS/MS. In parallel, the same analysis was performed using cells transfected with empty vector to exclude nonspecific binding. Proteins reproducibly detected across three independent experiments were extracted, and semiquantitative analysis based on identification frequency (IF)^51^ identified 196 proteins with IF > 0.01. Among those, 13 endosome-localized proteins were selected using GO Term and UniProt subcellular localization annotations (Fig. 1A). This group included the lysosomal protein TMEM55B and its binding partners, v-ATPase subunit Atp6v0d1 and Lamtor1 (Fig. 1A)^43^. Notably, Lamtor1 and TMEM55B are lysosome-localized proteins associated with cholesterol metabolism, suggesting a potential role in lipolysis-mediated cholesterol processing. Since TMEM55B regulates the v-ATPase complex, we focused on TMEM55B as a possible upstream regulator of PDZD8-mediated lysosomal function. Immunoprecipitation and immunoblot analysis confirmed the interaction between PDZD8 and TMEM55B (Fig. 1B).

**Figure 1.**
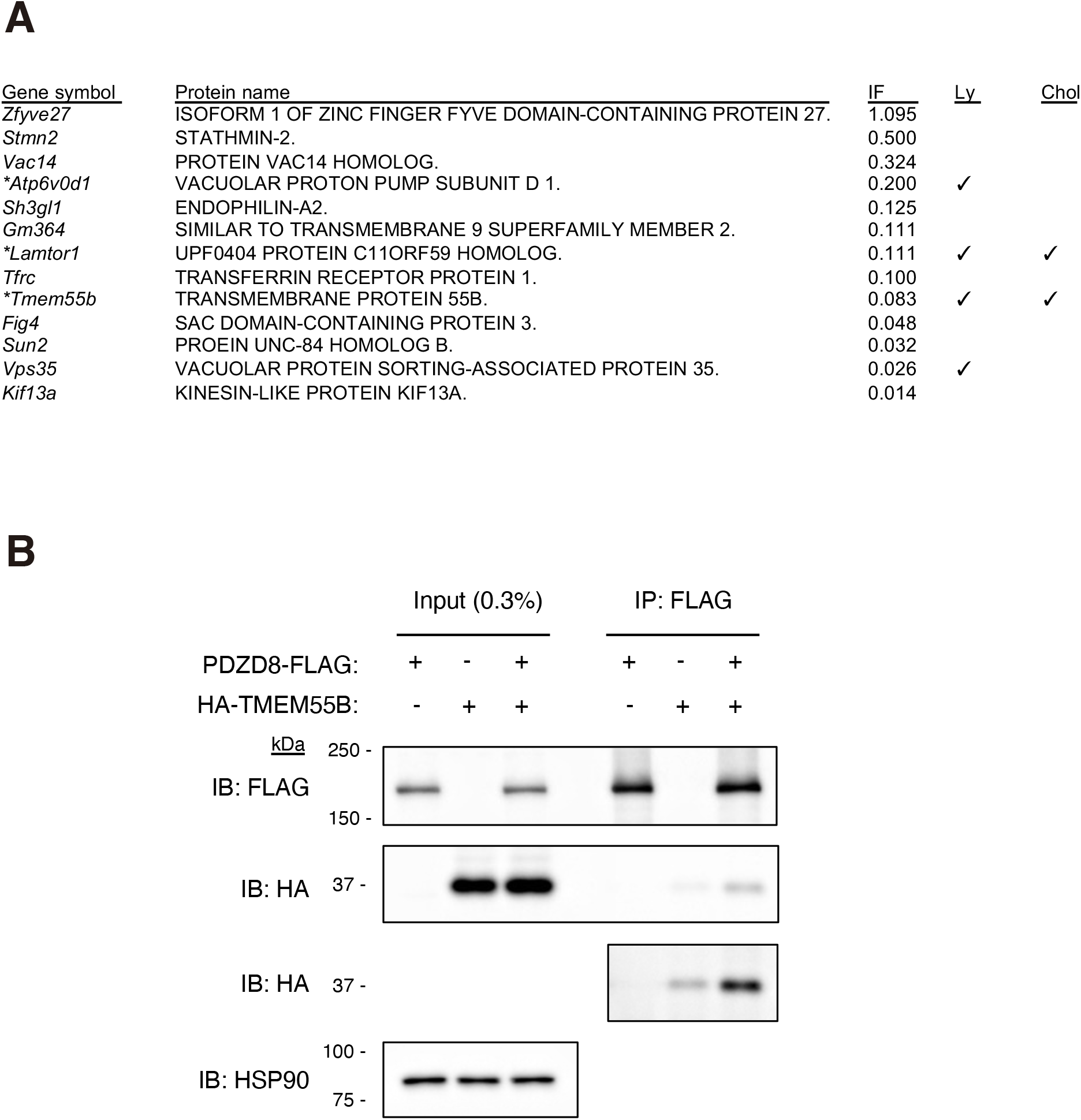
Identification of PDZD8-associated proteins by proteomic analysis. (**A**) Endosome-related proteins identified as PDZD8-associated proteins. Protein abundance was estimated semiquantitatively based on identification frequency (IF). Asterisks indicate proteins previously reported to associated with TMEM55B. Lysosome (Ly)-localized and cholesterol (Chol)-related proteins are indicated. Proteins were classified according to Gene Ontology (GO) terms and UniProtKB annotations. (**B**) HeLa cells were transfected with (+) or without (−) PDZD8-FLAG together with HA-TMEM55B. Cell lysates were subjected to immunoprecipitation-immunoblot (IP-IB) analysis using anti-FLAG, anti-HA, and anti-HSP90 antibodies. Input (0.3%) and FLAG-immunoprecipitated fractions were analyzed. Anti-HA immunoblots were additionally visualized using high-sensitivity detection reagents.

### TMEM55B is not essential for ER–LE/Ly tethering

Because PDZD8 functions as a tether at ER–LE/Ly MCSs, we examined whether its binding partner TMEM55B also contributes to this process. TMEM55B expression was suppressed in HeLa cells using siRNA (siTMEM55B), resulting in a marked reduction of both RNA and protein levels compared with control cells (siControl) (Fig. 2A, B). To assess ER–LE/Ly contacts, we employed a split-GFP system in which GFP fluorescence is reconstituted only when ER and lysosomal membranes are closely apposed^16,52^. Cells were co-transfected with ERj1-GFP(1–10) (ER-localized fragment) and LAMP1-GFP(11) (lysosomal fragment), together with mCherry to normalize expression levels. The GFP/mCherry fluorescence ratio per cell, reflecting ER–LE/Ly MCS formation, was quantified (Fig. 2C). No significant difference in GFP/mCherry ratio was observed between siTMEM55B and siControl cells (Fig. 2D). These results demonstrate that, unlike PDZD8, TMEM55B is not required for tethering of ER–LE/Ly MCSs.

**Figure 2.**
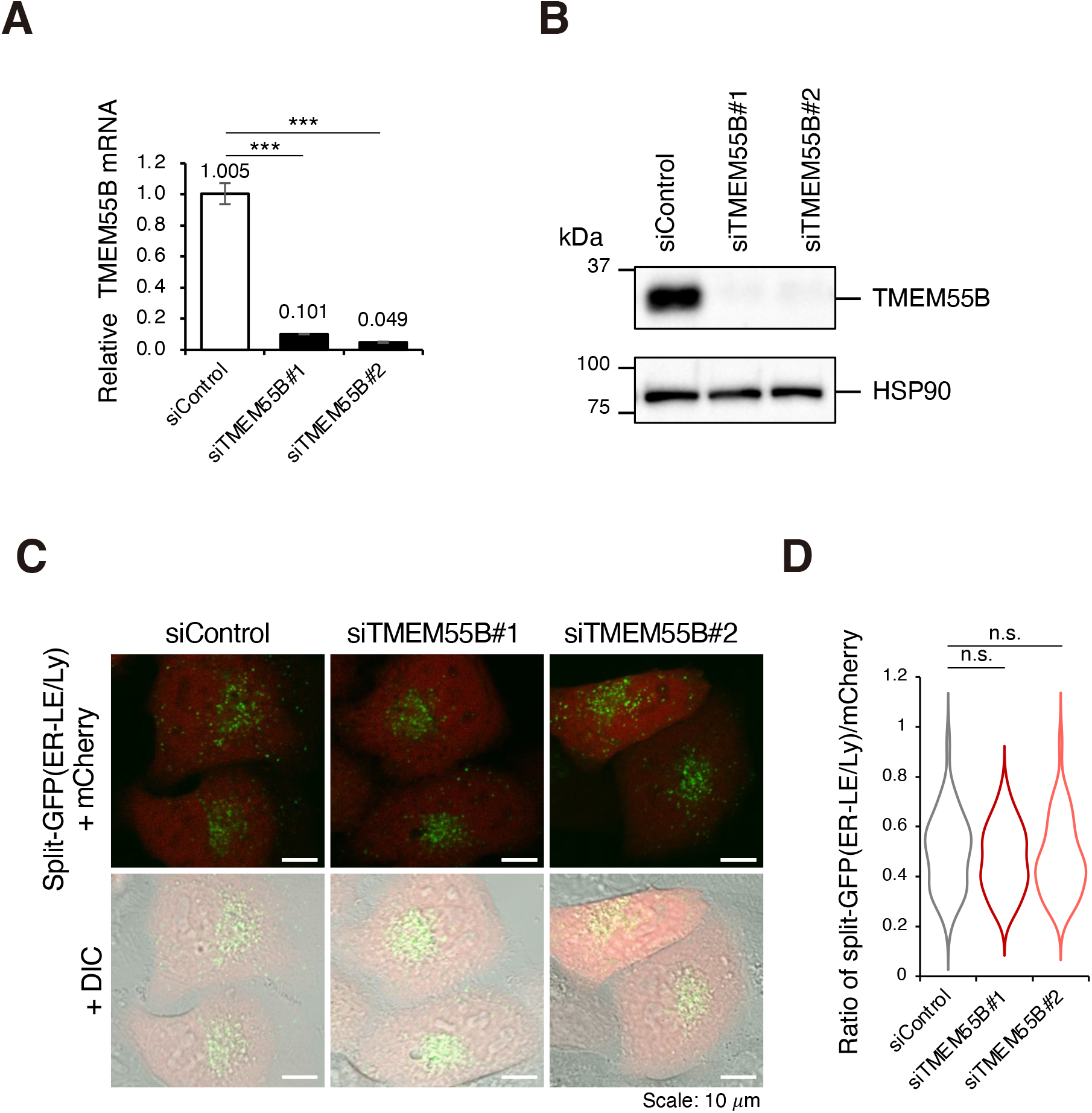
TMEM55B depletion does not affect ER–lysosome membrane contact formation. (**A, B**) HeLa cells were transfected with siControl, siTMEM55B#1, or siTMEM55B#2. TMEM55B mRNA levels were quantified by qRT–PCR (A), and protein levels were analyzed by immunoblotting with anti-TMEM55B and anti-HSP90 antibodies (B). Data are shown as mean ± SE (n = 9 for qRT–PCR). ***p < 0.001 (one-way ANOVA with Dunnett’s multiple-comparisons test). (**C**) HeLa cells were transfected with siControl, siTMEM55B#1, or siTMEM55B#2 together with split-GFP vectors ERj1–GFP(1–10) and LAMP1–GFP(11), as well as mCherry as a transfection control. Scale bar, 10 µm. (**D**) Quantification of split-GFP fluorescence intensity normalized to mCherry fluorescence in (C). n = 38, 36, and 37 cells for siControl, siTMEM55B#1, and siTMEM55B#2, respectively. n.s., not significant (Kruskal–Wallis test with Steel’s multiple comparisons test).

### TMEM55B regulates lysosomal acidification and activity

TMEM55B interacts with the v-ATPase complex and has been implicated in its regulation. We therefore hypothesized that TMEM55B contributes to lysosomal acidification and activity via v-ATPase. To test this, HeLa cells transfected with siControl or siTMEM55B were stained with LysoTracker (Fig. 3A) or LysoSensor (Fig. 3C), both of which increase in fluorescence intensity as lysosomal pH decreases. In both assays, siTMEM55B cells showed significantly reduced fluorescence compared with controls, indicating impaired lysosomal acidification (Fig. 3B, D). To assess the functional consequences in neurons, we suppressed TMEM55B expression in PC12 cells (Fig. 3E) and evaluated lipolysis using EGFP-PLIN2 as a lipid droplet (LD) marker and LysoTracker Red as a lysosomal marker (Fig. 3F). TMEM55B knockdown cells exhibited enlarged LDs (Fig. 3G) with increased LD fluorescence intensity per cell (Fig. 3H), as well as reduced LysoTracker fluorescence (Fig. 3I), consistent with defective lysosomal acidification and impaired lipolysis. Furthermore, the spatial association between lipid droplets and lysosomes, assessed by Pearson’s correlation analysis of EGFP-PLIN2 and LysoTracker signals, was significantly reduced in siTMEM55B cells compared with controls (Fig. 3J). Together, these results demonstrate that TMEM55B promotes lysosomal acidification and activity, thereby facilitating LD degradation and lipolysis.

**Figure 3.**
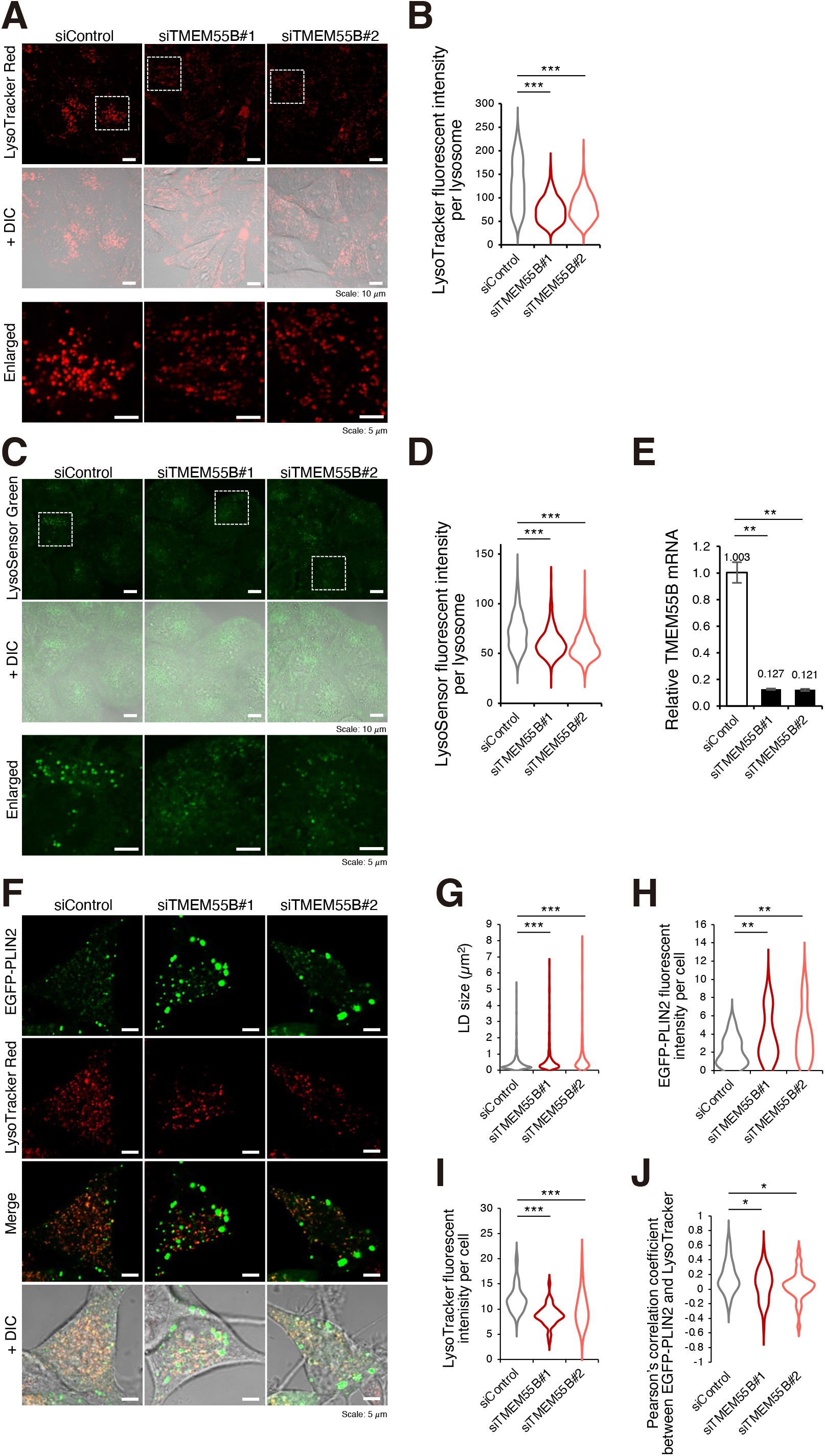
TMEM55B depletion impairs lysosomal acidification and lipolysis. (**A, C**) HeLa cells transfected with siControl, siTMEM55B#1, or siTMEM55B#2 were stained with LysoTracker Red (A) or LysoSensor Green (C) and analyzed by confocal microscopy. Boxed regions in the upper panels are shown at higher magnification below. Scale bars, 10 µm (upper and middle panels) and 5 µm (lower panels). (**B, D**) Quantification of LysoTracker Red (B) or LysoSensor Green (D) fluorescence intensity per lysosome in (A) (n = 746, 497, and 452 lysosomes for siControl, siTMEM55B#1, and siTMEM55B#2, respectively) or (C) (n = 703, 847, and 794 lysosomes, respectively). (**E**) qRT-PCR analysis of TMEM55B mRNA in PC12 cells transfected with siControl, siTMEM55B#1, or siTMEM55B#2. Data are shown as mean ± SE (n = 6). (**F**) PC12 cells were transfected with siControl, siTMEM55B#1, or siTMEM55B#2 together with EGFP-PLIN2 and stained with LysoTracker Red. (**G**) Quantification of lipid droplet (LD) area represented by EGFP-PLIN2-positive puncta in (F) (n = 627, 450, and 317 LDs for siControl, siTMEM55B#1, and siTMEM55B#2, respectively). (**H, I**) Quantification of mean EGFP-PLIN2 fluorescence intensity (H) and LysoTracker Red fluorescence intensity (I) per cell in (F) (n = 52, 41, and 35 cells for siControl, siTMEM55B#1, and siTMEM55B#2, respectively). (**J**) Pearson’s correlation coefficient between EGFP-PLIN2 and LysoTracker Red fluorescent signals in (F). ***p < 0.001, **p < 0.01 (Kruskal–Wallis test with Steel’s multiple comparisons test in B, D, and G–I); **p < 0.01, *p < 0.05 (one-way ANOVA with Dunnett’s multiple comparisons test in E and J).

### TMEM55B promotes Ca^2+^ release from the ER

Intracellular Ca^2+^ elevation promotes lysosome fusion with other organelles^44^, suggesting that lipolysis mediated by the PDZD8–TMEM55B complex could involve Ca^2+^ signaling at ER–LE/Ly MCSs. Since the ER is the major intracellular Ca^2+^ store, we investigated whether TMEM55B regulates Ca^2+^ release or uptake from the ER. HeLa cells were transfected with the ER-targeted Ca^2+^ indicator G-CEPIA1er^53^, which binds ER luminal Ca^2+^ and emits GFP fluorescence, and colocalization with the ER marker KDEL was confirmed by confocal microscopy (Fig. 4A, B). To assess ER Ca^2+^ dynamics, siControl or siTMEM55B cells were stimulated with ATP to induce transient ER Ca^2+^ release, followed by Ca^2+^ reuptake to the ER. In siTMEM55B cells, ATP-induced ER Ca^2+^ responses were markedly attenuated in siTMEM55B cells compared with controls (Fig. 4C). Quantitative analysis demonstrated that TMEM55B depletion reduced the peak ER Ca^2+^ response (Fig. 4D), decreased the initial rate of ATP-induced ER Ca^2+^ release (Fig. 4E), delayed the time required to reach peak release (Fig. 4F), and slowed the subsequent recovery phase (Fig. 4G). These findings indicate that TMEM55B facilitates ATP-stimulated transient Ca^2+^ release from the ER.

**Figure 4.**
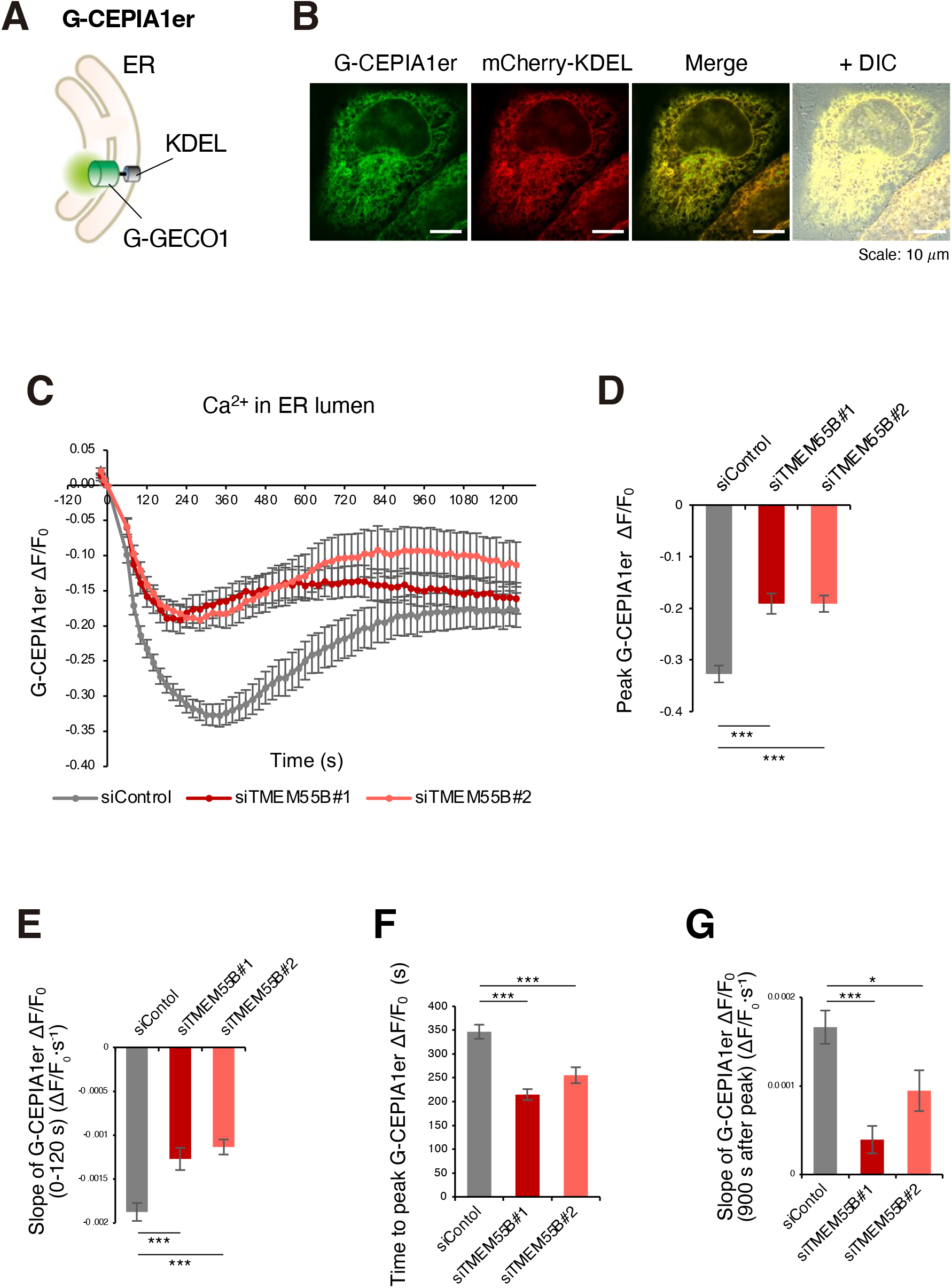
TMEM55B depletion impairs ER Ca^2+^ release. (**A**) Schematic illustration of G-CEPIA1er, an ER luminal Ca^2+^ indicator containing the G-GECO1 domain. (**B**) Confocal images of HeLa cells co-expressing G-CEPIA1er (green) and the ER marker mCherry-KDEL (red). Scale bar, 10 µm. (**C**) Time lapse analysis of G-CEPIA1er fluorescence (ΔF/F_o_) in cells transfected with siControl (gray, n = 30 fields), siTMEM55B#1 (red, n = 30), or siTMEM55B#2 (pink, n = 30). ATP (200 µM) was added at t = 0 s; Data are mean ± SE. (**D**) Quantification of peak ΔF/F_o_ values in (C). (**E**) Quantification of the slope of G-CEPIA1er ΔF/F_o_ during 0–120 s after ATP addition (ΔF/F_o_·s^−1^). (**F**) Quantification of the initial slope of G-CEPIA1er ΔF/F_o_ during the first 120 s after ATP addition (ΔF/F_o_·s^−1^). (**G**) Quantification of the slope of G-CEPIA1er ΔF/F_o_ during 900 s period following the peak (ΔF/F_o_·s^−1^). Data are mean ± SE. ***p < 0.001, *p < 0.05 (Kruskal–Wallis test with Steel’s multiple comparisons test in D, F, and G); ***p < 0.001 (one-way ANOVA with Dunnett’s multiple comparisons test in E).

### TMEM55B promotes Ca^2+^ release from lysosomes

We next investigated whether TMEM55B regulates Ca^2+^ release from lysosomes and how this affects ER Ca^2+^ dynamics. Lysosomal Ca^2+^ release and uptake were monitored using Ly-GG, which binds cytosolic Ca^2+^ at the lysosomal membrane and emits GFP fluorescence^54^ (Fig. 5A). Co-expression of Ly-GG with the lysosomal marker LAMP1 confirmed colocalization (Fig. 5B). ATP stimulation of siControl HeLa cells induced transient Ca^2+^ release from lysosomes. ATP-induced lysosomal Ca^2+^ responses were markedly attenuated in siTMEM55B cells compared with controls (Fig. 5C). Quantitative analysis revealed reduced rates of lysosomal Ca^2+^ release and reuptake (Fig. 5D, E). As a control, we examined mitochondrial Ca^2+^ using CEPIA3mt^53^, which binds mitochondrial matrix Ca^2+^, together with the mitochondrial marker Su9-RFP^55^ (Fig. 5F), confirmed colocalization (Fig. 5G). ATP stimulation induced transient mitochondrial Ca^2+^ increases as well as decrease, which were not significantly different between siControl and siTMEM55B cells (Fig. 5H–J). These results indicate that TMEM55B specifically regulates lysosomal Ca^2+^ dynamics without affecting mitochondrial Ca^2+^.

**Figure 5.**
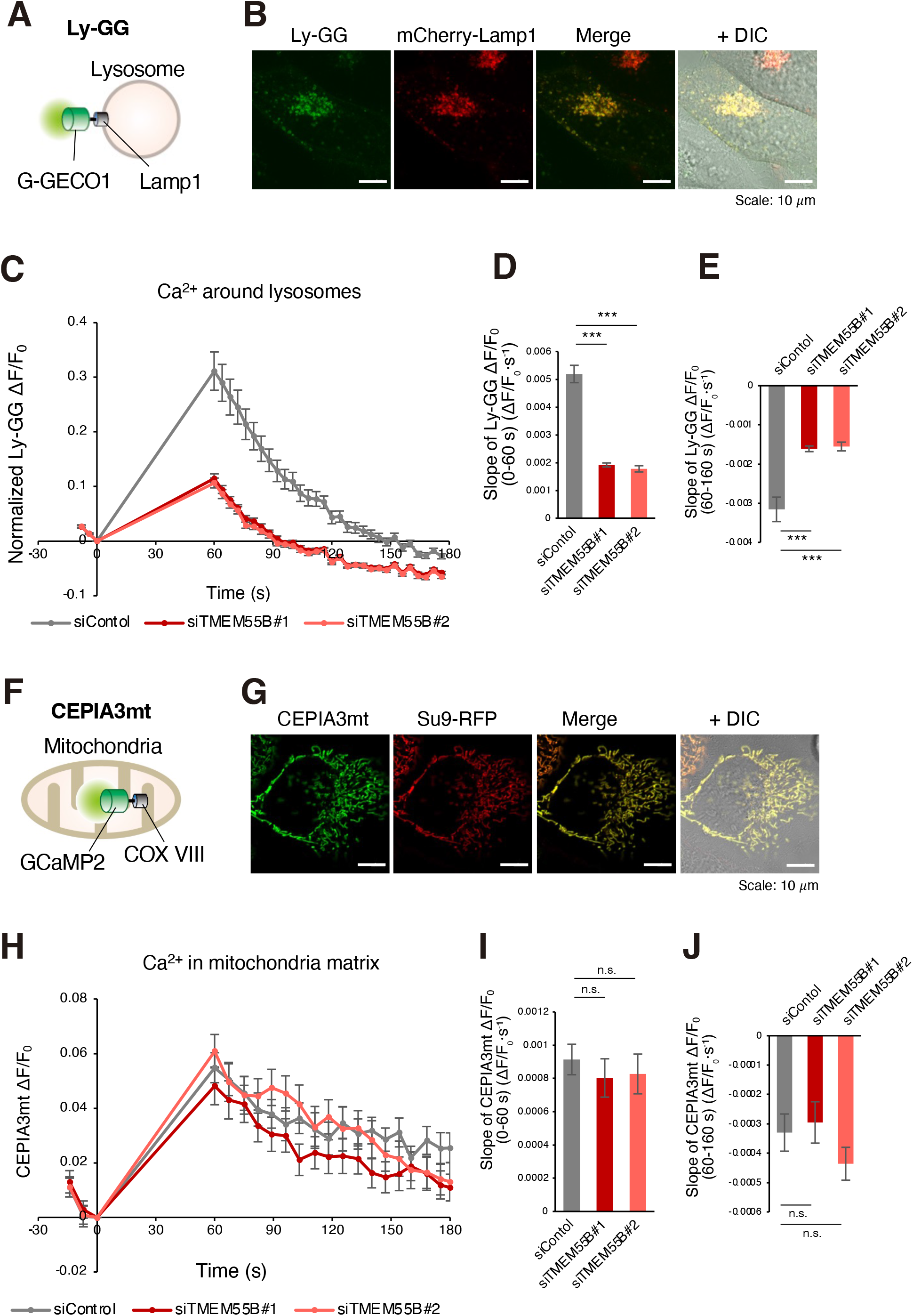
TMEM55B depletion impairs lysosomal Ca^2+^ dynamics. (**A**) Schematic illustration of Ly-GG, a lysosome-targeted Ca^2+^ indicator containing the G-GECO1 domain exposed to the cytosol. (**B**) Confocal images of HeLa cells co-expressing Ly-GG (green) and the lysosomal marker mCherry-LAMP1 (red). Scale bar, 10 µm. (**C**) Time lapse analysis of Ly-GG fluorescence (ΔF/F_0_) in cells transfected with siControl (gray, n = 29 fields), siTMEM55B#1 (red, n = 29), or siTMEM55B#2 (pink, n = 26). ATP (200 µM) was added at t = 0 s; Data are shown as mean ± SE and were corrected by subtracting signals from ATP-untreated cells to compensate for photobleaching. (**D, E**) Quantification of Ly-GG fluorescence slopes during 0–60 s (D) and 60–160 s (E) after ATP stimulation (ΔF/F_0_·s^−1^). (**F**) Schematic illustration of CEPIA3mt, a mitochondrial matrix-targeted Ca^2+^ indicator containing the GCaMP2 domain. (**G**) Confocal images of HeLa cells co-expressing CEPIA3mt (green) and the mitochondrial marker Su9–RFP (red). Scale bar, 10 µm. (**H**) Time lapse analysis of CEPIA3mt fluorescence (ΔF/F_o_) in cells transfected with siControl (gray, n = 25 fields), siTMEM55B#1 (red, n = 26), or siTMEM55B#2 (pink, n = 24). ATP (200 µM) was added at t = 0 s; Data are mean ± SE. (**I, J**) Quantification of CEPIA3mt fluorescence slopes during 0–60 s (I) and 60–160 s (J) after ATP stimulation (ΔF/F_0_·s^−1^). Data are mean ± SE. ***p < 0.001; (Kruskal–Wallis test with Steel’s multiple comparisons test in D and E); n.s., not significant (one-way ANOVA with Dunnett’s multiple comparisons test in I and J).

## Discussion

Lysosomes serve as central hubs for cellular metabolism, coordinating degradation, nutrient recycling, and signaling. Proper lysosomal acidification and calcium homeostasis are essential for lipid metabolism and neuronal function, and their dysregulation has been increasingly linked to neurodegenerative diseases. Despite this, the molecular mechanisms connecting lysosomal function to lipid regulation remain incompletely understood.

In this study, we identify TMEM55B as a key lysosomal regulator that interacts with the ER protein PDZD8 to control lysosomal acidification and calcium dynamics. TMEM55B regulates lysosomal acidification, Ca^2+^ dynamics, and lipid droplet turnover, revealing a mechanism linking lysosomal activity to lipid metabolism. These findings provide new insights into cellular lipid homeostasis and suggest potential targets for disorders associated with lysosomal dysfunction and cholesterol metabolism.

The lipid transfer protein PDZD8 mediates tethering of ER–LE/Ly MCSs and facilitates cholesterol metabolism through lipolysis via endosomal maturation. However, the mechanism by which PDZD8 promotes endosome maturation has remained unclear. Through complex analysis in neuronal cells, we identified Lamtor1 and TMEM55B as lysosome-localized, cholesterol-associated proteins among LE/Ly candidates. Given that TMEM55B regulates the v-ATPase complex, we focused on its intracellular functions.

We found that TMEM55B contributes to lysosomal acidification and activation. Loss of TMEM55B impaired lysosomal Ca^2+^ release and uptake, as well as ATP-stimulated Ca^2+^ release from the ER, while mitochondrial Ca^2+^ dynamics remained unaffected. In our system, lysosomal Ca^2+^ release peaked within 180 s after ATP stimulation (Fig. 5C), whereas ER Ca^2+^ release peaked later (Fig. 4C), consistent with calcium-induced calcium release (CICR) from the ER triggered by lysosomal Ca^2+^ release. These results indicate that TMEM55B promotes lysosomal acidification and Ca^2+^ dynamics and may facilitate CICR at ER–lysosome MCSs, thereby contributing to lipolysis (Fig. 6).

**Figure 6.**
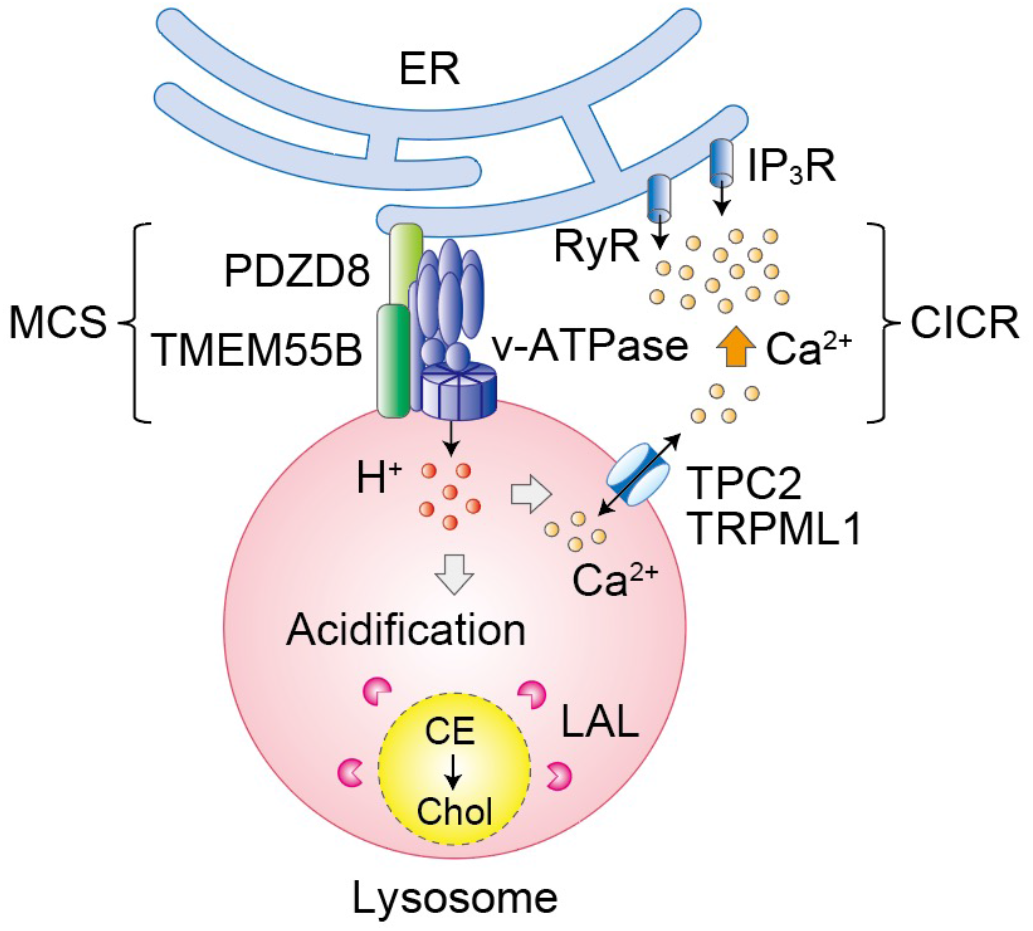
Proposed model by which TMEM55B promotes lipolysis through regulation of lysosomal acidification and Ca^2+^ dynamics at ER–lysosome membrane contact sites. The ER protein PDZD8 associates with the lysosomal protein TMEM55B, which promotes lysosomal acidification through regulation of v-ATPase complex. Enhanced lysosomal acidification facilitates the activity of lysosomal acid lipases, thereby promoting lipolysis and the conversion of cholesteryl esters (CEs) into free cholesterol. Acidification also enhances lysosomal Ca^2+^ signaling mediated by TPC2 and TRPML1, leading to Ca^2+^-induced Ca^2+^ release (CICR) through ER-localized RyR and IP3R channels at ER– lysosome membrane contact sites. This local Ca^2+^ signaling may further facilitate lysosome fusion with lipid droplets (LDs) or lipoprotein-containing endosomes, thereby promoting lipid degradation. ER, endoplasmic reticulum; MCS, membrane contact site; v-ATPase, vacuolar-ATPase; TPC2, two-pore channel 2; TRPML1, transient receptor potential mucolipin 1; RyR, ryanodine receptor; IP_3_R, inositol 1,4,5-trisphosphate receptor; CICR, Ca^2+^-induced Ca^2+^ release; LD, lipid droplet; LP, lipoprotein; CE, cholesteryl ester; Chol, cholesterol.

The PDZD8–TMEM55B complex therefore appears to contribute to multiple ER– lysosome MCS functions, including lysosomal regulation, Ca^2+^ dynamics, and cholesterol metabolism. Nevertheless, the effects of TMEM55B deficiency on lysosomal fusion and lipid droplet degradation were partial, failing to recapitulate the strong phenotype observed with PDZD8 deficiency. This suggests that additional, as-yet-unidentified mechanisms contribute to PDZD8-mediated lipolysis.

In summary, TMEM55B enhances lysosomal acidification via v-ATPase regulation, thereby increasing lysosomal enzyme activity and promoting lipolysis. Lysosomal acidification may additionally enhance lysosomal Ca^2+^ signaling by increasing the activity of lysosomal Ca^2+^ channels, thereby facilitating CICR at ER–lysosome MCSs. Whether TMEM55B-dependent CICR directly contributes to lipolysis remains to be determined.

## Methods

### Plasmids

EGFP and mCherry vectors were from Clontech (Mountain View, CA, US). Human cDNAs encoding PLIN2 (GenBank accession number NM_007408.4), LAMP1 (GenBank accession number BC025335), and KDEL with the signal sequence of calreticulin (GenBank accession number NM_004343) were amplified from HeLa cells by the polymerase chain reaction (PCR) with PrimeSTAR HS DNA polymerase (Takara, Shiga, Japan). pCMV-G-CEPIA1er (Addgene plasmid #58215) and pCMV-CEPIA3mt (Addgene plasmid #58219) were obtained from Addgene (Watertown, MA, US). Ly-GG was kindly provided by Dr. CW Taylor (University of Cambridge, UK). Vectors for split-GFP analysis, ERj1–GFP(1–10) and LAMP1–GFP(11), were kindly provided by Dr. Tamura (Yamagata University, Japan). The pSu9 cDNA was kindly provided by Dr. Mihara (Kyushu University, Japan). The mRFP cDNA was kindly provided by Dr. R.Y. Tsien (University of California, San Diego, US).

### Antibodies

Antibodies to TMEM55B were from Proteintech (Rosemont, IL, USA); antibodies to HSP90 were from BD Biosciences (Franklin Lakes, NJ, USA); antibodies to FLAG (mouse monoclonal M2 for immunoprecipitation and rabbit polyclonal for immunoblot analysis) were from Sigma-Aldrich (St. Louis, MO); antibody to HA (HA.11) was from Covance (Princeton, NJ).

### Cell culture, transfection, and retrovirus infection

HeLa cells (CCL-2, purchased from ATCC, US) were cultured under a humidified atmosphere of 5% CO_2_at 37°C in Dulbecco’s modified Eagle’s medium (Invitrogen Life Technologies) supplemented with 10% fetal bovine serum (Invitrogen Life Technologies). The culture medium for Plat-E cells was also supplemented with blasticidin (10 μg/ml). Cells were transfected with the use of XtremeGene9 reagents (Roche Diagnostics, Mannheim, Germany) according to the manufacturer’s protocol. For retroviral infection, Plat-E cells were transiently transfected with pMX-puro–based vectors and then cultured for 48 h. The retroviruses in the resulting culture supernatants were used to infect Neuro2A cells, and the cells were then subjected to selection with puromycin (1 μg/ml).

### Identification and categorization of PDZD8-associated proteins

Neuro2a cells infected with a retrovirus encoding FLAG–tagged mouse PDZD8 or with the corresponding empty virus were solubilized in a lysis buffer (40 mM HEPES-NaOH [pH 7.5], 150 mM NaCl, 10% glycerol, 0.5% Triton X-100, 1 mM Na_3_VO_4_, 25 mM NaF, aprotinin [10 μg/ml], leupeptin [10 μg/ml], 1 mM phenylmethylsulfonyl fluoride) to prepare whole-cell extracts. Lysates were cleared by centrifugation at 20,400 × g for 15 min at 4 °C. The clarified supernatants were subjected to affinity purification with anti-FLAG (M2)–agarose affinity gel (Sigma-Aldrich), and then proteins eluted with FLAG peptide (Sigma-Aldrich) was separated by SDS-PAGE and stained with silver. The stained gels were sliced into pieces, and the abundant proteins therein were subjected to in-gel digestion with trypsin. The resulting peptides were dried, dissolved in a mixture of 0.1% trifluoroacetic acid and 2% acetonitrile, and then applied to a nanoflow LC system (Advanced UHPLC; Michrom BioResources, Auburn, CA, USA) equipped with an L-column (C18, 0.15 by 50 mm, particle size of 3 μm; CERI, Tokyo, Japan). LC–MS/MS analysis was performed with an Orbitrap Velos Pro instrument (Thermo Fisher Scientific). All MS/MS spectra were compared with protein sequences in the International Protein Index (IPI, European Bioinformatics Institute) mouse version 3.44 with the use of the MASCOT algorithm. Assigned high-scoring peptide sequences were considered for correct identification. Identified peptides from 3 independent experiments were integrated and regrouped by IPI accession number. Proteins detected in ≥2 of 3 independent experiments from FLAG-PDZD8–expressing cells, but absent from control cells, were considered PDZD8-associated. Semi-quantitative abundance was estimated from identification frequency (IF; number of identified peptides per protein normalized to the number of theoretically detectable tryptic peptides). Functional categorization of PDZD8-associated proteins was categorized with use of the UniProtKB database (https://www.uniprot.org) based on Gene Ontology (GO) annotations. The proteins listed in figure 1a shows endosome-related (GO:0005768), as well as columns for lysosome (GO:0005764) or cholesterol-related as well.

### RNA interference, reverse transcription, and real-time PCR analysis

Stealth siRNAs targeted to human TMEM55B (siTMEM55B#1, 5’-ACGUGGAAGGCAAGAUGCAUCAGCA-3’; siTMEM55B#2, 5’-CAAGAUGCAUCAGCAUGUAGUCAAA-3’), rat TMEM55B (siTMEM55B#1, 5’-UGCUGAUGCAUCUUGCCUUCCACGU-3’; siTMEM55B#2, 5’-CAAGAUGCAUCAGCAUGUAGUCAAA-3’), and non-targeting control were obtained from Invitrogen Life Technologies. (Invitrogen Life Technologies, Carlsbad, CA, USA). Human siRNAs were introduced into HeLa cells with the use of Lipofectamine RNAiMAX (Invitrogen Life Technologies), and rat siRNAs were introduced into PC12 cells with the use of a Cell Line Nucleofector Kit V (VCA-1003) and Nucleofector instrument (Lonza, Basel, Switzerland) according to the manufacturer’s protocol. The cells were then cultured for 48 h (for HeLa) or 72 h (for PC12). Total RNA was isolated from cells with the use of an RNeasy Mini Kit (Qiagen, Hilden, Germany) according to the manufacturer’s protocol and was subjected to reverse transcription (RT) with the use of a QuantiTect Rev Transcription Kit (Qiagen). The resulting cDNA was subjected to real-time PCR analysis with Power SYBR Green PCR Master Mix in an ABI-Prism 7000 sequence detection system (Applied Biosystems, Foster City, CA). The amounts of each target mRNA were calculated and normalized by that of GAPDH mRNA. Primers for PCR (forward and reverse, respectively) were as follows: 5’-GTCAGTGGTGGACCTGACCTG-3’ and 5’-AAAGTGGTCGTTGAGGGCAAT-3’ for human GAPDH; 5’-CGAGTCTGCCAATCTCTCATC-3’ and 5’-AGTGCGGTCTGTGAACTCTGT-3’ for human TMEM55B; 5’-AAGGCTGAGAATGGGAAGCTG-3’ and 5’-GGAGATGATGACCCTTTTGGC-3’ for rat GAPDH; 5’-CTAGCCCGGATAGTGGGAGTG-3’ and 5’-TTCCTGGAGGTGCATTCTTGA-3’ for rat TMEM55B.

### Fluorescence imaging of live cells

LysoTracker Red DND-99 (0.2 µM) or LysoSensor Green DND-189 (0.2 µM) (Thermo Fisher Scientific, Rockford, IL, USA) were added to cells for 1 h at 37 °C according to the manufacturer’s protocol. HeLa and PC12 cells expressing plasmid-encoded fluorescent proteins for 24 h and 96 h respectively, or metabolically labeled with fluorescent probes were observed with an LSM800 confocal microscope (Zeiss, Oberkochen, Germany), and the images were processed for calculation of fluorescence intensity with ZEN imaging software (Zeiss) as well as Fiji (ImageJ, https://fiji.sc).

### Ca^2+^ imaging

Cell media of HeLa cells expressing organelle-targeted Ca^2+^ indicators were pre-incubated with HEPES-buffered saline (HBS; 137 mM NaCl, 5.9 mM KCl, 2.2 mM CaCl_2_, 1.2 mM MgCl_2_, 10 mM HEPES, 14 mM glucose, pH 7.4) for 1 h at 37 °C and then stimulated with ATP (200 µM) (Wako, Japan). The images were captured by Operetta high-content screening system (PerkinElmer, Waltham, MA, USA) using a ×20 objective (λ_ex 460–490 nm; λ_em 500–550 nm) at the time point of ATP addition (t = 0 s) and continued time points (t = X s) at a rate of one frame per 20 sec for total 1240 sec (G-CEPIA) or 180 sec (Ly-GG and CEPIA3mt). The concentration of Ca^2+^ for each time points was calculated by ΔF/F_0_ = (F_x_− F_0_)/F_0_. Images were analyzed by Harmony software (PerkinElmer).

### Immunoblot analysis

HeLa cells were lysed in Triton lysis buffer containing 40 mM HEPES-NaOH (pH 7.6), 150 mM NaCl, 0.5% Triton X-100, 10 mM MgCl_2_, 1 mM Na_3_VO_4_, 25 mM NaF, 1 mM EDTA, 1 mM PMSF, leupeptin (10 µg/mL), and 10% glycerol. After 10 min on ice, lysates were centrifuged at 15,000 × g for 10 min at 4 °C, and protein concentrations were determined with the Pierce BCA Protein Assay (Thermo Fisher Scientific). Extracts were resolved by SDS–PAGE, transferred to Immobilon-P membranes (Merck Millipore, Darmstadt, Germany), and probed with primary antibodies. Signals were developed with SuperSignal reagents (Thermo Fisher Scientific) and imaged on an LAS-4000 (GE Healthcare, Waukesha, WI, USA).

### Immunoprecipitation and immunoblot analysis

Protein extracts were subjected to immunoprecipitation for 1 h at 4°C with primary antibodies and protein G–Sepharose 4 Fast Flow (Amersham Biosciences, Uppsala, Sweden). The immunoprecipitates were washed three times with cell lysis buffer and then subjected to immunoblot analysis.

### Statistical analysis

Statistical analyses were performed with BellCurve for Excel software (Social Survey Research Information, Tokyo, Japan). Normality of data was assessed with the Shapiro– Wilk test. If the normality assumption was not met, the Kruskal–Wallis test with the Steel’s multiple comparisons test were applied. If data were normally distributed, One-way repeated-measures analysis of variance (ANOVA) with the Dunnett’s multiple comparisons test were performed. Quantitative data are presented as mean ± SE. Statistical significance of a *P* value is indicated as 0.01 ≤ *p* < 0.05 (*); 0.001 ≤ *p* < 0.01 (**); *p* < 0.001 (***); p ≥ 0.05 (n.s.). Sample size (*n*) are in the figure legends.

## Acknowledgements and Funding Information

We thank Y. Tamura for split-GFP cDNAs; E. Taylor for Ly-GG cDNA; K. Mihara for Su9 cDNA; K. I. Nakayama and T. Ohnishi for proteome analysis; K. Sato and K. Kato for Ca^2+^ imaging analysis; and the Research Equipment Sharing Center at Nagoya City University for assistance. This work was supported in part by KAKENHI grants from Japan Society for the Promotion of Science (JSPS) to M.S. (24K02122) and N.M. (JPMJSP2130).

## Author contributions statement

M.S. designed experiments, supervised the study, analyzed the data, and wrote the manuscript. N.M. and H.M. performed experiments.

## Additional information

### Data availability

The MS data have been deposited in the JPOST Repository Database as follows:

URL: https://repository.jpostdb.org

JPID and PXID are as follows;

Mock-FLAG complex in Neuro2A; (1) JPST004039,

PXD067776; (2) JPST004040, PXD067777; (3) JPST004041, PXD067779.

PDZD8-FLAG complex in Neuro2A; (1) JPST004042, PXD067780; (2) JPST004043,

PXD067782; (3) JPST004044, PXD067783.

### Supplementary datasets

Raw data are provided within the supplementary information file.

